# Significant increase in root exudation of 2′-deoxymugineic acid (DMA) as a response to zinc deficiency in rice

**DOI:** 10.64898/2026.03.16.711580

**Authors:** Claudia Rocco, Gerald Larrouy-Maumus, Matthias Wissuwa, Colin Turnbull, Ramon Vilar, Dominik Weiss

## Abstract

- Zinc (Zn) deficiency limits rice productivity and poses a risk to human health, particularly in populations reliant on rice-based diets. Although rice germplasm exhibits wide variation in Zn-deficiency tolerance, the underlying physiological mechanisms remain poorly resolved. Evidence across the literature for Zn-deficiency–induced secretion of 2′-deoxymugineic acid (DMA) is inconsistent. This study clarifies the role of DMA secretion as a Zn-deficiency stress response.
- We developed and validated a sensitive LC–ESI–Q–TOF–MS method for selective detection of DMA in rice root exudates. Five rice genotypes with contrasting Zn-deficiency tolerance were grown hydroponically and DMA secretion measured.
- Zn-deficiency increased DMA exudation across all genotypes, with sensitive genotypes also showing higher secretion compared with control, supporting DMA’s role as a general response to Zn stress rather than being restricted to efficient genotypes.
- Fold-change responses exceeded previous studies, likely due to more severe stress exposure. Our results confirm that DMA secretion is induced under Zn-deficiency in rice as part of the micronutrient stress response. However, the lack of increased Zn uptake indicates that additional tolerance mechanisms are involved. These findings reconcile inconsistencies in the literature and position DMA secretion as an important, but not exclusive, component of Zn-deficiency adaptation in rice.

## 2 Introduction

Zinc (Zn) is an essential micronutrient for all living organisms (Wairich, 2022). Adequate zinc concentrations in crop are essential for human health and agricultural productivity. In crop, zinc is necessary as cofactor for enzymes, transcription factors and proteins structures (Maret, 2009, Ozyildirim and Baltaci, 2023). Currently, about 49% of agricultural soils worldwide are deficient in Zn (Graham, 2008, FAO, 2022), leading to reduced yield and low nutritional quality of plant production, particularly cereals. Rice (*Oryza sativa* L.) is a staple food that contributes 35 to 59 % of the daily intake for approximately three billion people (Selby-Pham et al., 2017, Wairich, 2022). Deficiency of zinc is a common occurrence in rice, causing mild to severe deficiency in more than two billion people worldwide representing a major threat to human health, particularly in countries where nutrient-deficient rice is the major source of calorie intake (WHO, 2007, Hanikenne and Bouché, 2023). In humans, zinc plays several essential roles, including the development and function of immune cells, neuronal function (e.g. synaptic plasticity), growth and development, and reproductive health (Singh et al., 2016, Lee et al., 2020, Kim and Lee, 2021). Pregnant women and children are particularly affected by zinc deficiency, which can lead to premature birth and premature death (Nakandalage et al., 2016, Selby-Pham et al., 2017). People that rely on rice as staple food can suffer from what is known as “hidden hunger” when trace metals, including zinc, are present at insufficient levels (Mushtaq et al., 2024), a problem that has become a global issue (U.N.E.P., 2021).

Micronutrient deficiencies are often caused by the insolubility or low solubility of trace metal nutrients due to particular soil conditions such as organic matter content > 3 wt%, neutral or alkaline pH, high electrical conductivity (EC), high carbonate content and low redox potential due to prolonged waterlogging, preventing their dissolution into the bioavailable pool (Alloway, 2008, Khush et al., 2012). Rice germplasm has a large variation in tolerance to Zn-deficiency (Wissuwa et al., 2006), but the mechanisms of its tolerance, such as enhanced root growth, Zn uptake and internal transport efficiency, and root exudation of organic compounds, remain unclear hindering efforts to breed Zn-efficient rice varieties with high grain Zn content (Mori et al., 2015).

It is widely known that to promote the uptake of poorly bioavailable Fe(III), graminaceous plants, including rice, have developed the ability to exude low molecular weight organic ligands, called siderophores (Schenkeveld et al., 2014, Suzuki et al., 2021). Once exuded, siderophores mobilize the poorly soluble Fe(III) from soil forming high-affinity soluble complexes (Suzuki et al., 2021, Yamagata et al., 2022). 2′-deoxymugineic acid (DMA) is the main α-hydroxycarboxylate phytosiderophore, that is exuded by rice plants during Fe(III) deficiency stress (Dell’Mour et al., 2010). Rice roots secrete DMA in the rhizosphere using specific transporter (TOM) to chelate the insoluble Fe(III) ion from soil and take up DMA-Fe complexes through yellow stripe 1-like (YSL) transporters (Tsednee et al., 2012, Suzuki et al., 2021, Yamagata et al., 2022) (Fig. 1).

Numerous studies (Wissuwa et al., 2006, Suzuki et al., 2008, Wissuwa et al., 2008, Arnold et al., 2010, Widodo et al., 2010, Nishiyama et al., 2012, Markovič et al., 2017, Northover et al., 2021) have proposed that, under Zn-deficiency, tolerant rice cultivars may exude DMA able to bind the insoluble zinc(II) from the soil matrix, following the same mechanism as Fe(III) chelation. Speciation modelling and isotopic analyses have shown a strong correlation between DMA secretion and Zn content, providing indirect evidence for a potential role of DMA in Zn(II) acquisition (Widodo et al., 2010, Weiss et al., 2021).

Despite existing studies provided circumstantial support for DMA excretion under Zn-deficiency, direct evidence of its secretion remains unclear, partly due to analytical challenges. Several methods have been applied for DMA detection, including hydrophilic interaction liquid chromatography–electrospray ionisation mass spectrometry (HILIC-ESI-MS) for direct separation and identification of DMA and its metal complexes, and derivatisation-based liquid chromatography–electrospray ionisation–time-of-flight mass spectrometry (LC–ESI– TOF-MS) using FMOC-Cl to enhance detectability (Xuan et al., 2006, Kakei et al., 2009, Tsednee et al., 2012). Nevertheless, sensitivity and specificity remain critical challenges due to its low abundance in root exudates and structural similarity to related phytosiderophores (Xuan et al., 2006, Selby-Pham et al., 2017). Targeted liquid chromatography–tandem mass spectrometry (LC-MS/MS) methods (Tsednee et al., 2012), have improved separation, identification, and quantification, allowing a more reliable assessment of DMA dynamics in purified and relatively abundant extracts. Nevertheless, these methods still have limitations for direct measurement in root exudates due to extremely low concentrations and the chemical complexity of the exudate matrix. The present study therefore combines optimised exudate collection and direct LC–MS/MS analysis, enabling sensitive and accurate quantification of DMA in unpurified root exudates. This approach allows quantitative assessment of DMA exudation directly from root exudate samples and provide a simple, direct, and highly sensitive analytical approach that addresses limitations reported for earlier methods.

This study aims to clarify the role of DMA secretion as a potential Zn-deficiency stress response. To this end, i) we first developed and validated a robust liquid chromatography– electrospray ionisation–quadrupole–time-of-flight mass spectrometry (LC–ESI–Q–TOF–MS) protocol for accurate identification and quantification of 2’-deoxymugineic acid (DMA) directly in rice root exudates under controlled hydroponic conditions; ii) we then tested the hypothesis that Zn deficiency stimulates DMA secretion in rice and assessed the variation in this response among genotypes with contrasting Zn-deficiency tolerance; iii) we finally perform a comparative analysis of DMA exudation and Zn-uptake trends across studies to contextualise our observations within previously published work and contribute to a clearer understanding of DMA’s role in Zn(II) acquisition.

To address these questions, five rice genotypes (A69-1, RIL46, IR55179, IR64 and IR74) with contrasting Zn-deficiency tolerance were grown under controlled hydroponic conditions. Hydroponic cultivation allowed us precise manipulation of zinc availability and facilitated direct collection of root exudates, which is difficult to achieve in soil due to sorption processes and microbial degradation. Moreover, hydroponics reduces environmental variability, enabling reliable comparisons among genotypes. While this system does not replicate soil conditions, it is well suited for mechanistic investigation of plant responses to nutrient deficiency. Root exudates were collected and analysed using a targeted LC–MS/MS method optimised for DMA detection and quantification. Following chromatographic separation, DMA was identified based on its characteristic mass-to-charge ratio and fragmentation pattern, providing high selectivity and minimizing interference from co-eluting metabolites. This approach was selected because DMA occurs at low concentrations in complex root exudate matrices, and LC–MS/MS allows sensitive, robust, and reproducible quantification while effectively discriminating DMA from structurally related compounds.

## 3 Materials and methods

### 3.1 Chemicals

All reagents were of analytical grade. Unless otherwise stated, all chemicals and reagents were purchased from Sigma-Aldrich or Fisher Scientific and used as received.

### 3.2 Plant materials

Five rice (*Oryza sativa* L.) genotypes belonging to the *indica* sub-species were selected for this study with seed being provided by the Japan International Research Center for Agricultural Sciences (JIRCAS). Their contrasting tolerance to Zn deficiency had been established in previous studies with A69-1, RIL46 and IR55179 being classified as tolerant to Zn deficiency, IR64 as moderately sensitive, and IR74 as highly sensitive (Frei et al., 2010, Mori et al., 2015).

### 3.3 Pot experiment and extraction of metabolites

All five rice genotypes were germinated for 4 days at 30 °C on paper soaked with distilled water. After germination, the uniform seedlings were transferred in 4-L black plastic pots and grown hydroponically in a growth chamber (day: 28°C, 13 h of light at µmol m^−2^ s^−1^; night: 11 h at 24 °C; humidity: 75%) without stirring or aeration. The first week plants were grown in in half-strength modified Yoshida solution (Yoshida et al. 1976) with the following composition at full strength: NH_4_NO_3_ 1.42 mmol L^−1^, KH_2_PO_4_•2H_2_O 0.05 mmol L^−1^, K_2_SO_4_ 0.5 mmol L^−1^, CaCl_2_•2H_2_O 1 mmol L^−1^, MgSO_4_•7H_2_O 1 mmol L^−1^, MnCl_2_•4H_2_O 9 μmol L^−1^, (NH_4_)_6_Mo_7_O_24_•4H_2_O 0.07 μmol L^−1^, H_3_BO_3_ 18.5 μmol L^−1^, CuSO_4_•5H_2_O 0.16 μmol L^−1^, Fe(EDTA) 36 μmol L^−1^, ZnSO_4_•7H_2_O 0.15 μmol L^−1^. The pH of the nutrient solution was 4.7 and renewed every 2-3 days. An aliquot of nutrient solutions prepared for the experiment was collected and analysed for zinc and iron content by using Agilent 8900 Triple Quadrupole ICP-MS to monitor potential contamination. Unless otherwise stated, no Zn and Fe reagents were added to the respective deficient nutrient solution. The detected concentration of Fe and Zn unintentionally added to the solution up to 0.20 and 0.09 μM respectively in the full-strength nutrient solutions (Table S1). The plants for the deficient treatment were always supplied with lower concentration of Fe(III) and Zn(II) salts respectively. During the last week of experiments, Fe(III)-EDTA and ZnSO_4_ were omitted for the respective deficiency treatments. The root exudates collected, and the plants were harvested after 4 weeks from the germination. The root exudates were stored in the freezer at -80 ºC. Frozen 50 ml aliquots of exudate solution were freeze-dried at -80 ºC for 5 days and re-dissolved in 1 ml sterilized MilliQ water (resistivity ∼18MΩcm). Exudates were then analysed for DMA content.

All experiments were conducted in a randomized complete block design with three replicates.

### 3.4 Collection of root exudates and HPLC analysis

For exudate collection, the roots 18 plants of each genotype from each replicate were rinsed with deionized water and separately soaked in 200 ml of MilliQ water (resistivity ∼18MΩcm) 1 h before illumination and covered with aluminium foil to prevent light degradation of exudates. Root exudates were collected for 4 h. The antimicrobial agent Micropur (Katadyn Products Inc., Wallisellen, Switzerland) was added to the water to prevent microbial degradation of the DMA after collection. After the sampling, exudates were filtered through a 0.20 µm syringe filter into 50 ml plastic tubes and immediately frozen at -80°C. Condensed and microfiltered samples were subjected to HPLC analysis.

### 3.5 LC MS-MS determination of DMA in the root exudates

Frozen 50 ml aliquots of exudate solution were freeze-dried at -80 ºC for 5 days and re-dissolved in 1 ml sterilized MilliQ water. Exudates were then analysed for DMA content.

The data were acquired with an Agilent 1290 Infinity II HPLC coupled to a 6545 LC/Q TOF system. Chromatographic separation was performed with an Agilent InfinityLab Poroshell 120 HILIC-Z (2.1 × 100 mm, 2.7 μm (p/n 675775-924)) column. The HILIC-Z methodology was optimized for polar acidic metabolites. For easy and consistent mobile-phase preparation, a concentrated 10x solution consisting of 100 mM ammonium acetate (pH 9.0) in water was prepared to produce mobile phases A and B. Mobile phase A consisted of 10 mM ammonium acetate in water (pH 9) with a 5-μM Agilent InfinityLab Deactivator Additive (p/n 5191-4506), and mobile phase B consisted of 1.0 mM ammonium acetate (pH 9) in 10:90 (v:v) water/acetonitrile with a 5-μM Agilent InfinityLab Deactivator Additive (p/n 5191-4506). The following gradient was applied at a flow rate of 0.5 ml/min: 0 minutes, 100% B; 0-11.5 minutes, 70% B; 11.5-15 minutes, 100% B; 12-15 minutes, 100% B; and 5-minutes of re-equilibration at 100% B. Accurate mass spectrometry was performed using an Agilent Accurate Mass 6545 QTOF apparatus. Dynamic mass axis calibration was achieved by continuous infusion after the chromatography of a reference mass solution using an isocratic pump connected to an ESI ionization source operated in negative-ion mode. The following parameters were used: sheath gas temperature, 300 °C; nebulizer pressure, 40 psig; sheath gas flow, 12 l min^-1^; capillary voltage, 3000 V; nozzle voltage, 0 V; and fragmentor voltage, 115 V. The data were collected in centroid 4 GHz (extended dynamic range) mode. The detected *m/z* data were deemed to represent metabolites, which were identified based on unique accurate mass-retention times and MS/MS fragmentation identifiers for masses exhibiting the expected distribution of accompanying isotopomers. The typical variation in the abundance of most of the metabolites remained between 5 and 10% under these experimental conditions.

Considering that a large proportion of plant root exudates comprised high-molecular-weight compounds such as proteins and mucilage (Chai and Schachtman, 2022), the DMA results were normalized by measuring in each individual samples the residual protein content of the plant exudates using the BCA assay kit (Thermo®).

### 3.6 Data analysis

All data are presented as mean values with relative standard deviation of three replicates and were subjected to an analysis of variance (ANOVA) using the Duncan test (P ≤ 0.05) as a post hoc test for the separation of the means determined using SPSS version 17.0 (SPSS Chicago, Illinois). Changes in Zn accumulation were evaluated by comparing the ratios of Zn concentrations in plants grown under Zn-sufficient and Zn-deficient conditions. Statistical analysis of Zn uptake was performed using a two-way ANOVA, with environmental treatment (control vs. Zn deficiency) and genotype (IR74 and RIL46) as fixed factors.

## 4 Results and discussion

### 4.1 Detection and identification of DMA in the root exudates

For the quantification and identification of DMA, we used HPLC-ESI-Q-TOF-MS in negative ionization mode. This ionisation mode is preferred for acidic compounds like DMA since they ionize more efficiently and produce higher signal intensity and lower background noise compared to positive ion mode, resulting in more robust identification and quantification in complex biological samples (Dell’mour et al., 2012, Liigand et al., 2017). In ESI(-) mode, DMA (C_12_H_20_N_2_O_7_) was detected with major peaks at [M-H]^-^ 303.1198 *m*/*z*, as expected from previous work (Xuan et al., 2006). Fig. 2 shows examples of chromatograms detecting DMA in the root exudates of rice grown under different nutrient conditions (deficient and control). When rice plants were subjected to Fe-deficiency, secretion of DMA in IR55179 genotype increased four times more than the control and No-Zn treatment (Table S2). The increase abundance of DMA exudation in rice under iron deficiency condition is well documented in the literature (Bashir et al., 2010, Selby-Pham et al., 2017).

**Fig. 2.**
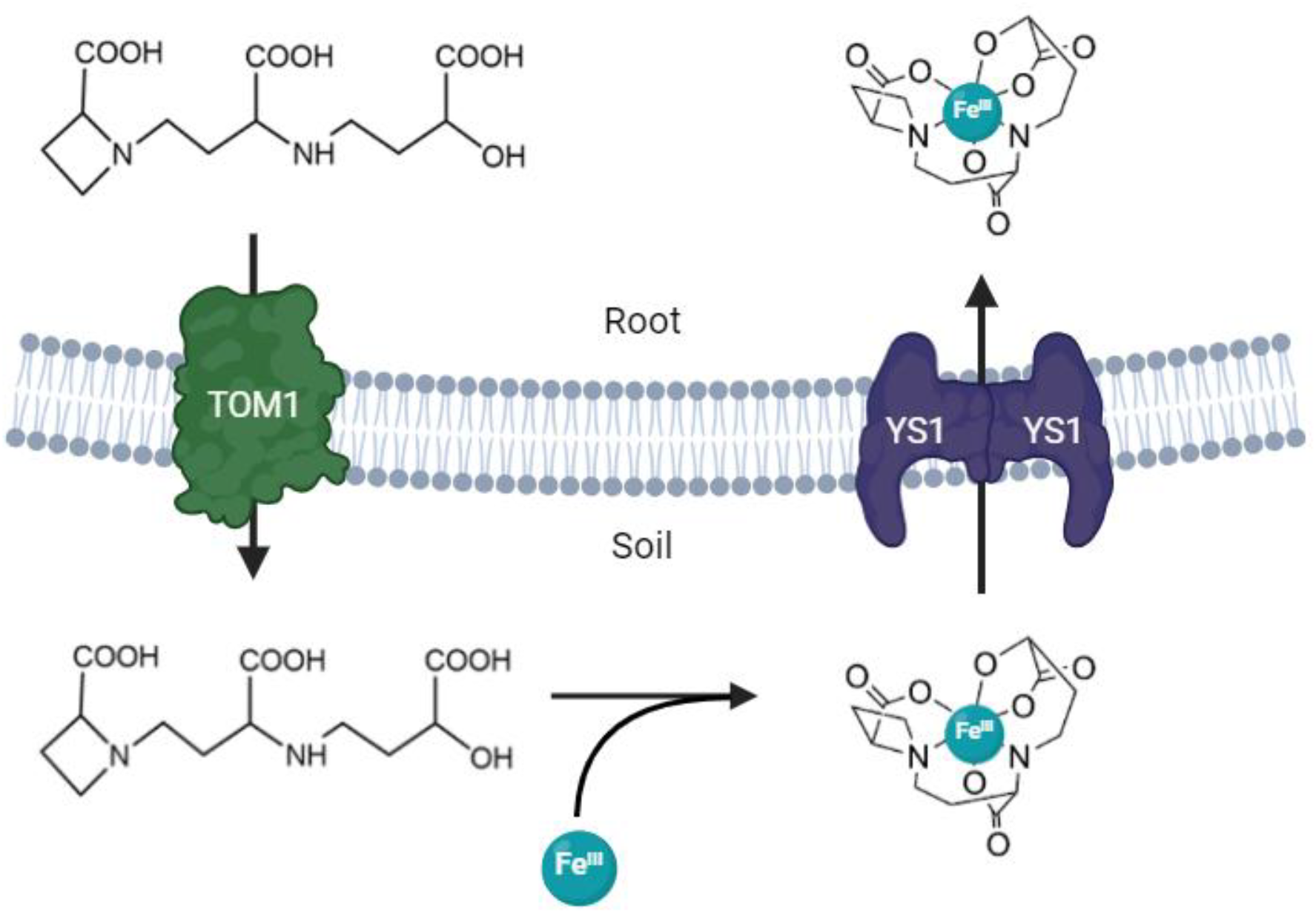
Iron uptake strategy in rice roots. Schematic representation of phytosiderophore-mediated iron acquisition in rice. Roots secrete 2′-deoxymugineic acid (DMA) into the rhizosphere via Transporters of Mugineic Acids (TOM). Fe(III)–DMA complexes are formed and subsequently taken up by root epidermal cells through Yellow Stripe 1 (YS1/YS1-like) membrane transporters.

**Fig. 2.**
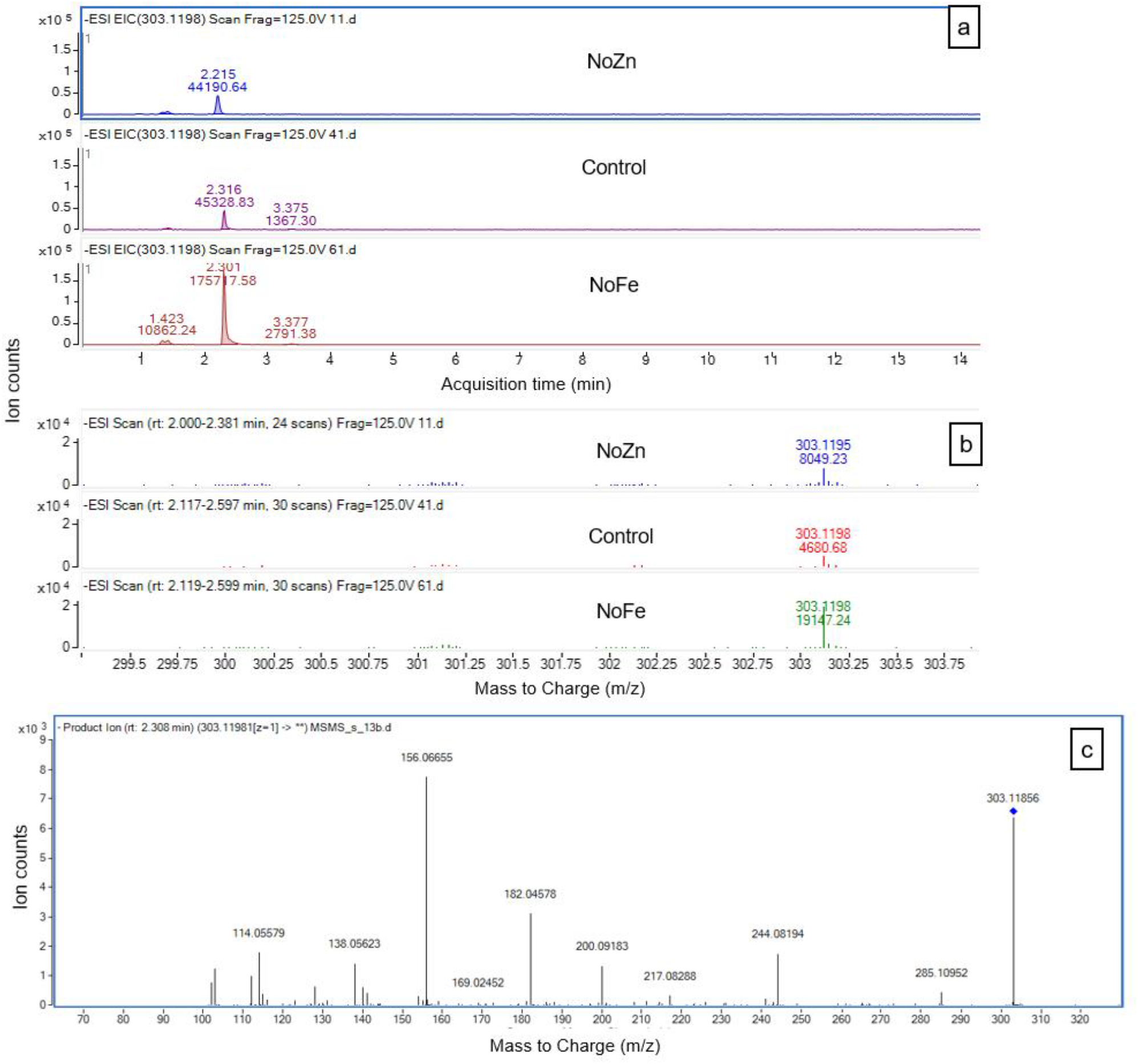
High performance liquid chromatography (HPLC)–ESI–QTOF–MS analysis of 2′-deoxymugineic acid (DMA) detected in rice root exudates under different nutrient deficiency conditions. (a) Extracted ion chromatograms (*m/z* 303.1198) showing treatment dependent DMA peak intensities. (b) Accurate mass spectra confirming DMA with detected precursor ions at *m/z* 303.11955 (NoZn), 303.11977 (Control) and 303.11988 (NoFe). (c) MS/MS fragmentation spectrum of the DMA precursor ion, displaying characteristic fragment ions that confirm the structural identity of DMA. Peaks and fragmentations were detected in ESI-mode using the calculated *m/z* of the DMA.

To validate that the peak in the chromatograms corresponded to the presence of DMA in the samples, we used the mass fragmentation pattern (Fig. 2b). DMA in the rice exudate samples was identified by its MS/MS spectrum (Fig. 2c). The calculated and observed *m*/*z* values of the identified compounds and their fragmentation patterns are provided in Table 1. The fragmentation observed during our study are comparable with those detected in previous work (Tsednee et al., 2012) using comparable LC-ESI-MS/MS to characterise DMA fragmentation from concentrated root exudate samples and provided in online metabolome database (hmdb.ca).

**Table 1.**
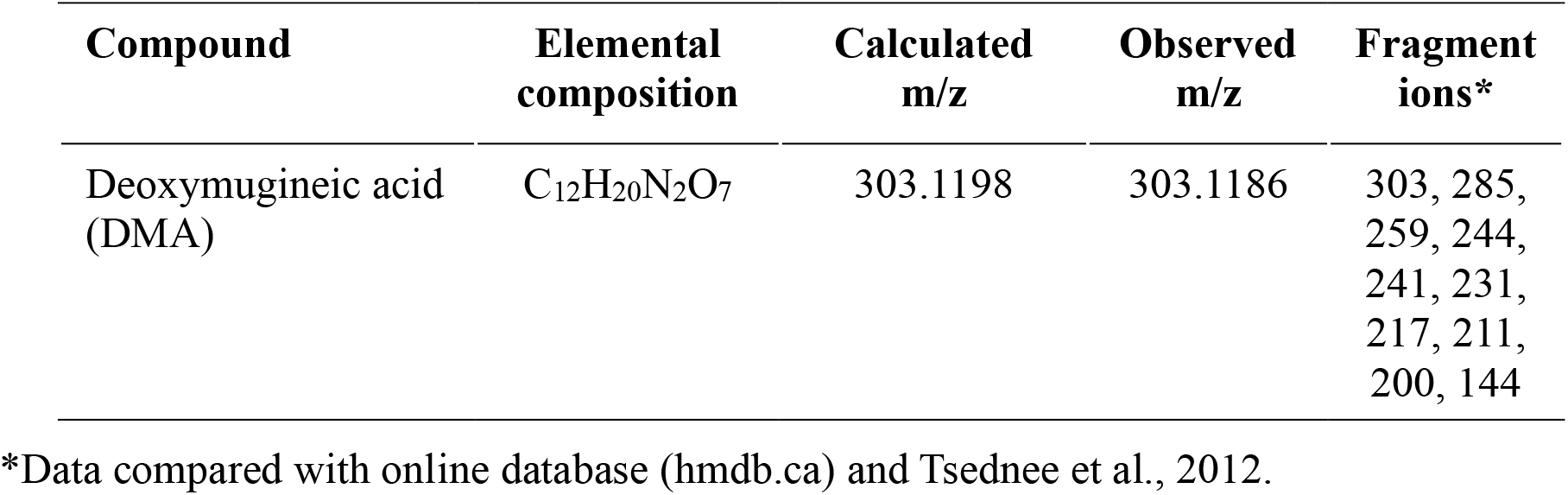
Calculated and observed *m/z* values of DMA in rice exudate as processed in ESI-mode QTOF.

| Compound | Elemental composition | Calculated $m/z$ | Observed $m/z$ | Fragment ions* |
| --- | --- | --- | --- | --- |
| Deoxymugineic acid (DMA) | $C_{12}H_{20}N_2O_7$ | 303.1198 | 303.1186 | 303, 285, 259, 244, 241, 231, 217, 211, 200, 144 |

\*Data compared with online database (hmdb.ca) and Tsednee et al., 2012.

The combination of optimised chromatographic separation, accurate mass detection in negative ion mode, and confirmation by MS/MS fragmentation provides a robust framework for the direct detection of DMA in root exudates. This supports the development of a simple, direct, and highly sensitive analytical method that overcomes the limitations of earlier indirect or derivatisation-based approaches (Suzuki et al., 2008, Widodo et al., 2010). Comparable strategies using LC–ESI–TOF–MS have demonstrated the advantages of direct detection for phytosiderophores such as DMA and nicotianamine, particularly in terms of sensitivity and sample throughput (Kakei et al., 2009). The use of HILIC-based methodologies has been shown to be suitable for the analysis of polar and acidic metabolites in plant systems, enabling improved resolution and reproducibility (Tsednee et al., 2012, Tang et al., 2016). In this context, our approach provides an important advance for the study of rice root exudation under nutrient deficiency stress, facilitating a reliable assessment of the dynamics of DMA secretion in comparison with previous methods.

### 4.2 Plant growth: Influence of nutrient deficiency on different rice genotypes

At harvest time, plants subjected to iron and zinc deficiency exhibited mild chlorosis, whereas overall growth and physiological activity were maintained (Fig. S1). We chose to grow rice plants for three weeks, as DMA exudation can be reliably detected in 21-day-old seedlings (Widodo et al., 2010) and LC–ESI–TOF/QTOF method is sensitive enough to quantify DMA from young plant material, confirming that early seedling stages provide suitable samples for direct analysis (Kakei et al., 2009). Extending growth further would risk additional stress due to the limited pot size used.

Fig. 3 and Fig. S2 illustrate the effects of Fe and Zn deficiencies on the growth (biomass and length) of five rice genotypes. The genotypes are classified based on their tolerance to Zn deficiency, with three Zn-tolerant lines (A69-1, RIL46, IR55179) and two Zn-sensitive lines (IR64, IR74).

**Fig. 3.**
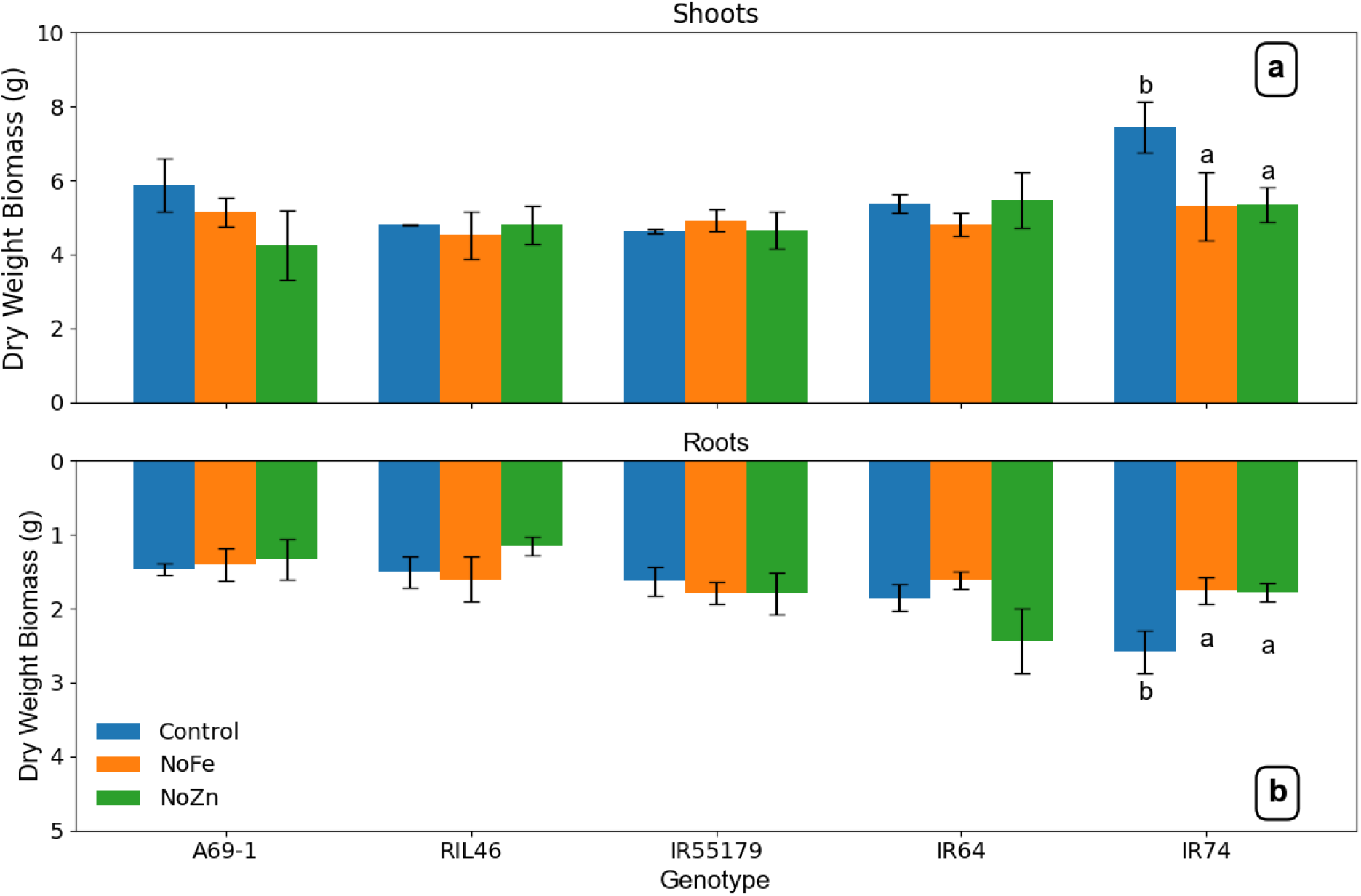
Mean dry weight (± SD, *n* = 18) of rice shoots (a) and roots (b) for five genotypes (A69-1, RIL46, IR55179, IR64, IR74) grown under three nutrient regimes: control (+Zn/+Fe), Fe deficiency (NoFe) and Zn deficiency (NoZn). Different letters indicate significant differences (*P* < 0.05).

In Fig. 3, the dry biomass of shoots and roots in genotypes A69-1, IR64, and IR74 was significantly higher in control plants compared to RIL46 and IR55179, indicating genotypic differences in growth potential under hydroponic conditions. Under Zn- and Fe-deficiency treatments, the sensitive genotype IR74 showed a significant reduction in shoot and root biomass of approximately 30% relative to the control, confirming its susceptibility to micronutrient stress. In contrast, IR55179 and RIL46 maintained their biomass across the treatments, showing no significant differences between control and deficiency conditions and confirming their tolerance. These results confirm the contrasting biomass responses of IR74 and RIL46 reported previously (Widodo et al., 2010, Rose et al., 2012), and extend these findings by including three additional genotypes.

Fig. 4 shows the effects of Fe and Zn deficiency on root-to-shoot biomass and length ratios. For biomass ratios (Fig. 4a), genotype A69-1 exhibited a significantly lower root/shoot ratio under control and Fe-deficient conditions, whereas the other genotypes maintained similarly high ratios. Zn deficiency elicited the strongest responses: the root/shoot ratio increased in A69-1 and IR64, remained unchanged in IR55179 and IR74, and decreased in RIL46. For length ratios (Fig. 4b), Zn deficiency also altered root elongation relative to shoots in a genotype-dependent manner, with A69-1 and IR64 showing proportionally longer roots, while RIL46 showed reduced root elongation. These results suggest that under Zn deficiency, rice preferentially allocates resources to root growth, but the magnitude and direction of this response vary among genotypes, reflecting contrasting strategies of nutrient foraging.

**Fig. 4.**
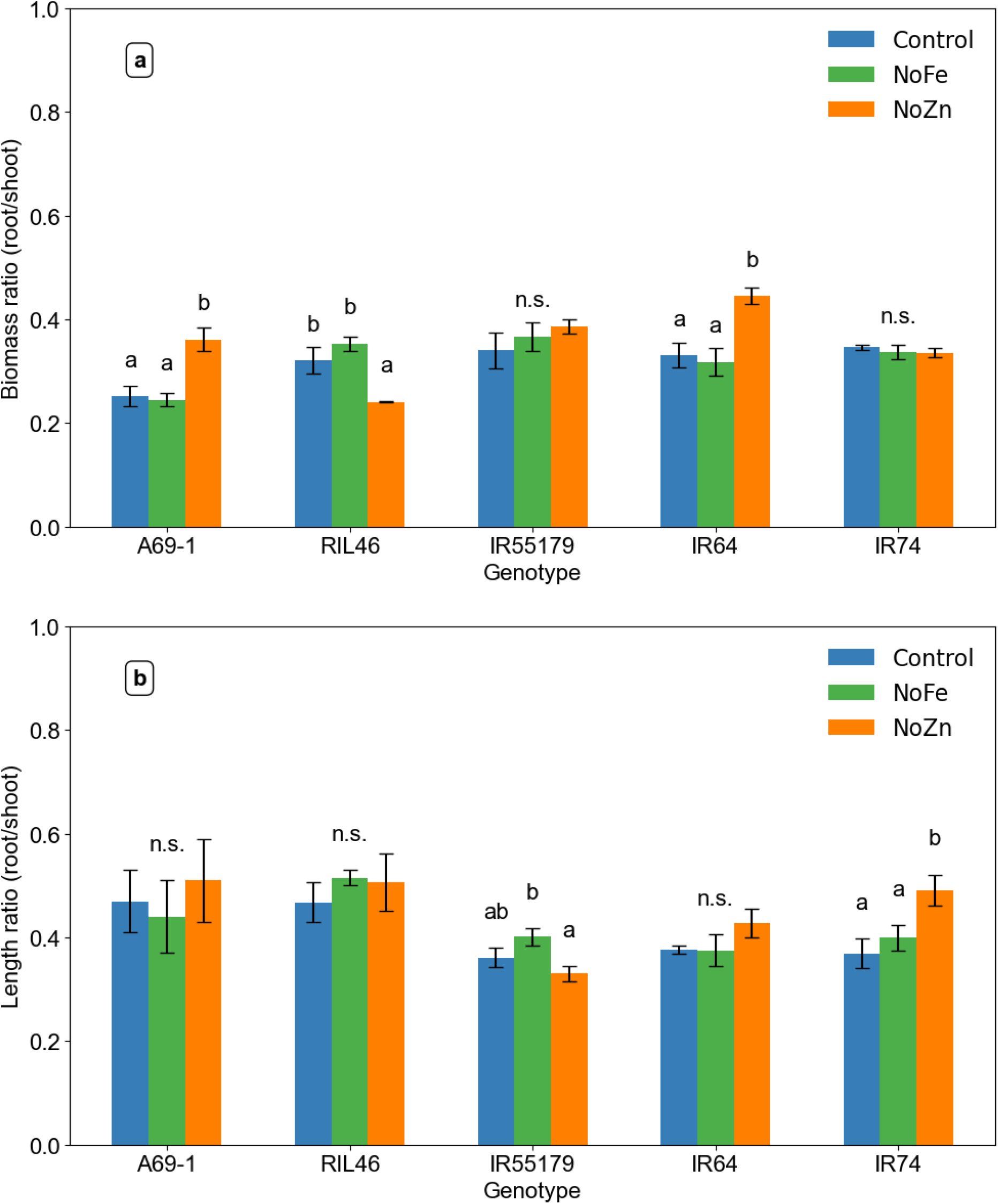
Mean ± SD (n = 18) root-to-shoot ratios for rice genotypes based on (a) dry biomass and (b) root and shoot length under full nutrient solution (Control), Fe deficiency (NoFe), and Zn deficiency (NoZn). Different letters above bars indicate statistically significant differences (P < 0.05); bars labelled “n.s.” indicate no significant difference

A69-1 and IR64 genotypes exhibit adaptive plasticity, ability of a plant to modify its growth or physiology in response to environmental conditions (Napier et al., 2022), by increasing root development to enhance nutrient uptake under Zn deficiency. This is consistent with other findings (Mori et al., 2015, Nanda and Wissuwa, 2016), reporting that genotypes tolerant to Zn-deficiency develop more extensive root systems under Zn stress compared to the sensitive genotypes. RIL46, despite being previously classified as Zn-tolerant, showed a decrease in root/shoot ratio and no significant change in biomass across treatments. This observation suggests that, under the hydroponic Zn-deficiency conditions tested here, RIL46 does not rely on root proliferation as a tolerance mechanism, which aligns with reports that its Zn-deficiency tolerance is instead associated with maintaining root growth (Widodo et al., 2010, Nanda and Wissuwa, 2016).

Root-to-shoot length ratio (Fig. 4b) show that under full nutrient solution (control), the tolerant genotypes A69-1 and RIL46 have a significantly high length ratio compared to IR55179, IR64, and IR74, indicating genotypic differences in growth potential. Under Fe- and Zn-deficient conditions, a similar trend is observed except for IR74 which significantly increased its length ratio under Zn deficiency condition compared to the control. Contrary, IR55179 significantly increase its ratio under Fe deficiency treatment but decreased it under Zn deficiency. The other genotypes showed no significant changes between the control and the deficiency treatments (Fig. S2). This result indicates that under Zn deficiency condition, IR74 produced longer roots without increasing root biomass, suggesting a morphological adjustment to explore a larger volume of the growth medium, a response commonly observed in nutrient-limited plants (Agathokleous et al., 2019).

### 4.3 DMA exudation from rice root increases under nutrient deficiency conditions

Fig. 5 shows the results of DMA root exudation by different rice genotypes in response to zinc and iron deficiency stress in a hydroponic experiment. Analysis of root exudates in the hydroponic solution revealed that DMA release differed significantly under nutrient-deficient conditions compared with the control. After 3 weeks from transplanting, DMA abundance per gram of root dry weight (DMA g^-1^ root d.w.) in the exudate solution was not significantly different between genotypes in the control treatment (Fig. S3), ranging from 245±205 to 548±248 DMA g^-1^ root. Fe-deficiency strongly induces DMA exudation in all genotypes. The strongest induction (57.7-fold) is observed in genotype A69-1, ranging from 3 to 18 times more compared to the other genotypes (Fig. S3), and the weakest (5.5-fold) in genotype RIL46. Values ranged from a minimum of 1770±228 DMA g^−1^ root d.w. for IR64 (moderate intolerant) to a maximum of 31235±5635 DMA g^−1^ root d.w. for A69-1 (tolerant).

**Fig. 5.**
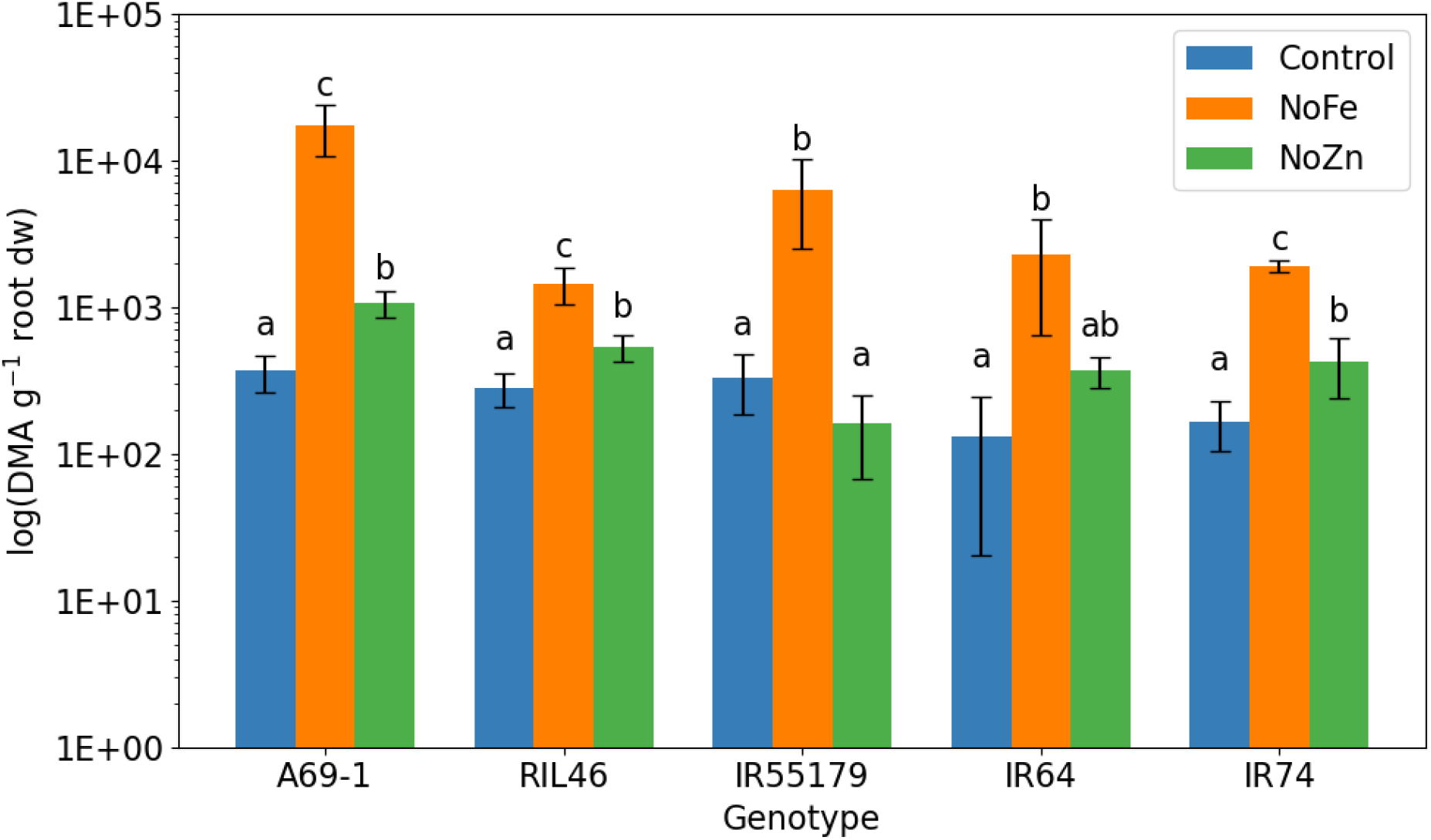
Deoxymugineic acid (DMA) exudation by rice roots per each genotype grown under Fe- and Zn-deficient conditions in hydroponic. DMA abundance is normalised to root dry weight (DMA g^−1^ root d.w.) and represents relative signal intensity detected by LC–ESI– Q-TOF–MS. Bars represent mean values ± SD. Different letters above bars indicate significant difference among treatments within each genotype (ANOVA, P < 0.01).

Zn deficiency induced DMA exudation in most, but not all, genotypes and to a smaller extent than Fe deficiency. Relative to their respective control, the strongest induction is observed in genotype A69-1(2.5-fold), and the weakest (0.5-fold) in genotype IR55179, this being the only genotype where DMA exudation is lower under Zn deficiency. This may indicate that DMA-driven Zn solubilization is not a tolerance mechanisms in this genotype, which agrees with observations of Lee and co-workers (2017) that tolerant IR55179 tolerated very low tissue Zn concentrations, exhibited lower accumulation of stress markers, and maintained more stable root function as base of its tolerance mechanisms (Lee et al., 2017).

Interestingly, the two sensitive genotypes IR64 and IR74, presented significant differences in DMA exudation between nutrient deficiency treatments and the control. Both genotypes exhibited a strong and similar response under Fe deficiency, with DMA exudation increasing 7.2-fold in IR64 and 7.9-fold in IR74 relative to the control. Under Zn deficiency, the increase was more moderate, with a 3.6-fold rise in IR64 and a 1.8-fold rise in IR74 compared to the control. Overall, the no Zn–induced fold change was smaller than under no Fe and broadly comparable among most genotypes, with the exception of IR55179, which showed a decrease. Although IR55179 has been described as a Zn-efficient genotype, this has mainly been observed in Zn-deficient soils, where increased root number and growth enhance Zn acquisition (Nanda and Wissuwa, 2016). In hydroponic systems, where rhizosphere-mediated soil interactions are absent, these differences may be less pronounced. The significantly higher exudation of DMA detected in NoZn-treated sensitive genotypes supports the hypothesis that DMA is involved in the response of Zn deficiency stress, even in sensitive genotypes. This is a key new finding, indicating that increased DMA secretion represents a general response to Zn deficiency rather than being restricted to Zn-efficient genotypes.

Under nutrient deficiency conditions, the tolerant genotype A69-1 shows the highest DMA exudation per gram of root (d.w.). Relative to its control treatment, DMA exudation is 57.7-fold higher under Fe-deficiency and 2.5-fold higher under Zn-deficiency (Fig. 5). Across genotypes, A69-1 secreted on average 11-fold more DMA under Fe-deficiency and 2.6-fold more DMA under Zn-deficiency than the other lines, representing the most tolerant genotype in both deficiency conditions. RIL46 showed a strong deficiency response, with DMA exudation increasing 5.5-fold under Fe deficiency and 1.5-fold under Zn deficiency compared with its control. Under Zn deficiency, DMA exudation was 1.8-fold higher in A69-1 and 0.8-fold higher in RIL46 than in the sensitive genotype IR74, consistent with previous findings. Overall, the greater DMA exudation in A69-1 and RIL46 under nutrient deficiency is consistent with their higher tolerance compared with IR64 and IR74.

The unit used in Fig. 5 and Fig. S3 (DMA g^−1^ root d.w.) represents the LC–ESI–Q-TOF–MS signal intensity normalised to root dry weight, which allows comparison of relative DMA secretion across genotypes within this experiment.

Our results suggest a role for DMA in the rice response to nutrient deficiency stress. All genotypes responded to iron deficiency with strongly increased DMA release, confirming a role of DMA in Fe complexation and uptake. The response to Zn deficiency was less pronounced, but all genotype except IR55179, shows 1.5 to 3.6 -fold higher DMA exudation than control. However, the magnitude of this response varied among genotypes. The tolerant lines A69-1 and RIL46 exhibited the highest absolute DMA exudation, particularly under Fe deficiency, where A69-1 released up to 18-fold more DMA than the other genotypes. Sensitive genotypes such as IR64 and IR74 also increased DMA secretion under stress, suggesting that DMA exudation is a general stress response though not sufficient alone to confer tolerance. These results suggest that while DMA secretion is commonly induced by nutrient deficiency, the absolute amount released contributes to genotypic differences in tolerance and represents a useful trait for breeding micronutrient-efficient rice. DMA secretion under Zn deficiency was 2 to 40-fold lower than under Fe-deficiency, confirming that DMA release is much more strongly stimulated by Fe-deficiency than by Zn-deficiency (Suzuki et al., 2008).

### 4.4 Comparison with existing studies on root exudation and zinc uptake

In table 2, we compare our exudation data with findings reported by Widodo et al. (2010), who measured DMA exudation in 21-day-old seedlings of rice genotypes RIL46 and IR74 grown hydroponically in Yoshida nutrient solution. For the Zn-deficiency treatment, Zn was omitted from the nutrient solution seven days prior to exudate collection, while the control treatment received 0.1 µM Zn.

**Table 2.**
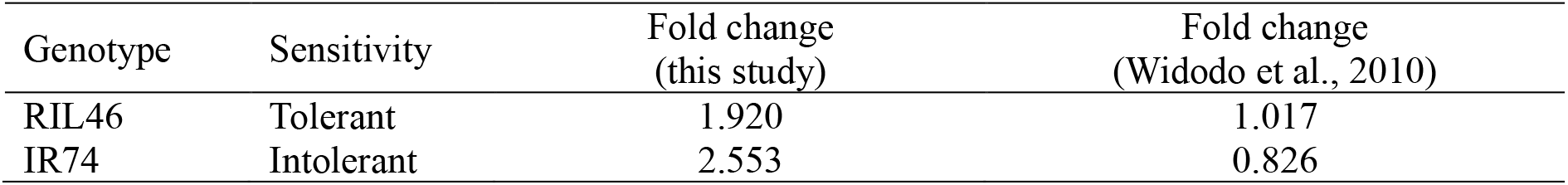
Relative changes in 2′-deoxymugineic acid (DMA) exudation for two rice genotypes (RIL46-tolerant; IR74-intolerant) grown under Zn-deficient and Zn-sufficient conditions in this study, compared with values reported by Widodo et al. (2010). Widodo et al. measured DMA exudation in 21-day-old rice seedlings grown hydroponically in Yoshida nutrient solution. Fold change was calculated as the ratio of DMA released under Zn-deficient conditions relative to Zn-sufficient controls.

| Genotype | Sensitivity | Fold change<br>(this study) | Fold change<br>(Widodo et al., 2010) |
| --- | --- | --- | --- |
| RIL46 | Tolerant | 1.920 | 1.017 |
| IR74 | Intolerant | 2.553 | 0.826 |

Zinc uptake values presented in Table 3 and Fig. 6 were obtained from previously published field and hydroponic studies (Arnold et al., 2010, Widodo et al., 2010, Rose et al., 2012, Impa et al., 2013). Field experiments by Widodo et al. (2009) and Arnold et al. (2010), performed in soils naturally deficient in Zn, involved Zn fertilisation, with plants harvested approximately four weeks after transplanting. Hydroponic experiments by Impa et al. (2013) and Rose et al. (2012) were performed in controlled greenhouse environments using Yoshida nutrient solution or agar-based media. Zn treatments were defined as Zn-deficient (0.15 µM) and Zn-sufficient (1.5 µM), and harvest times ranged from 14 to 21 days after treatment initiation. Total Zn uptake (µg plant^−1^) was evaluated by comparing values reported for Zn-deficient and Zn-sufficient conditions at similar early growth stages. Fold changes were calculated as the ratio of Zn-deficient to Zn-sufficient treatments. All data were obtained from previously published studies; we performed the comparative analysis and did not conduct any uptake experiments.

**Table 3.** Zinc accumulation (µg Zn plant^−1^) in two rice genotypes (RIL46-tolerant; IR74-intolerant) under zinc-sufficient (+Zn) and zinc-deficient (NoZn) conditions, based on data from Widodo et al. (2009), Arnold et al. (2010), and Rose et al. (2012). Fold change was calculated as the ratio of DMA released under Zn-deficient conditions relative to Zn-sufficient controls. Reduction represents the percentage decrease in Zn under Zn-deficient conditions relative to the corresponding Zn-sufficient (+Zn) treatment for each genotype and study.

| Genotype | Sensitivity | Treatment | Widodo et al.<br>2010 | Arnold et al.<br>2010 | Rose et al.<br>2012 |
| --- | --- | --- | --- | --- | --- |
|  |  |  | (µg Zn plant<sup>-1</sup>) |  |  |
| RIL46 | Tolerant | Control (+Zn) | 193 | 164.32 | 28 |
|  |  | NoZn | 31 | 26.09 | 14 |
| IR74 | Intolerant | Control (+Zn) | 212 | 188.89 | 40 |
|  |  | NoZn | 18 | 18.46 | 12 |
| Fold changes | RIL46 |  | 0.161 | 0.159 | 0.500 |
|  | IR74 |  | 0.085 | 0.098 | 0.300 |
| Reduction (%) | RIL46 |  | 84 | 84 | 50 |
|  | IR74 |  | 92 | 90 | 70 |

**Fig. 6.**
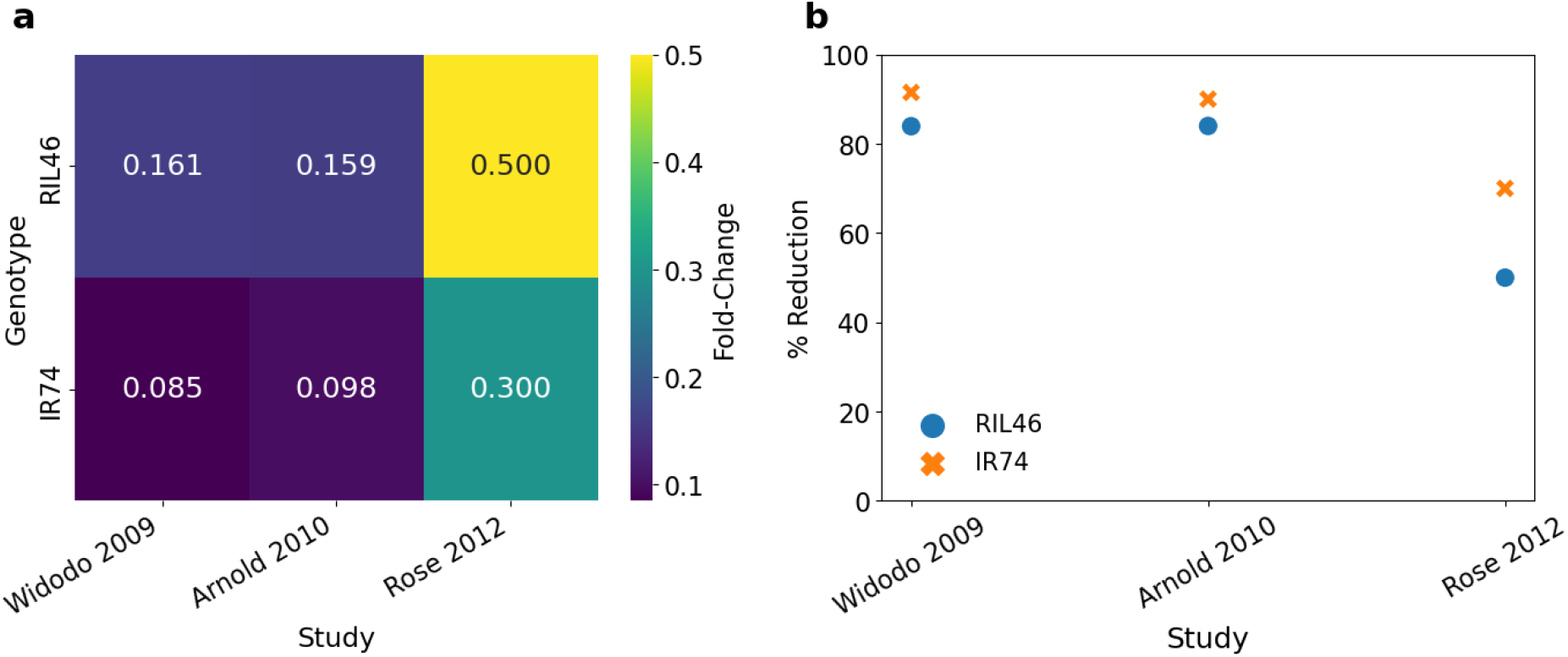
Comparative response of rice genotypes RIL46 (Zn-tolerant) and IR74 (Zn-intolerant) under Zn-sufficient (+Zn) and Zn-deficient (NoZn) conditions across multiple studies. (a) Heatmap showing fold-change ratios (NoZn/+Zn) for Zn accumulation. (b) Scatter plot illustrating percentage reduction in Zn content under deficiency. Lower fold-change and higher percentage reduction indicate greater sensitivity to Zn deficiency.

#### 4.4.1 Comparative evaluation of DMA exudation responses under Zn deficiency

Direct numerical comparison of DMA levels across studies is challenging due to methodological differences, but fold-change comparisons remain feasible.

Table 2 shows that DMA exudation under Zn deficiency increased 1.9-fold in RIL46 and 2.6-fold in IR74. These responses are stronger than those reported by Widodo et al. (2010), who observed fold-changes of 1.0 in RIL46 and 0.8 in IR74. This indicates a stronger DMA response to Zn deficiency in our experiment for both genotypes.

In Widodo et al. (2010), genotype differences were more pronounced, with RIL46 consistently exuding more DMA than IR74 under both Zn-sufficient and Zn-deficient conditions, suggesting strong genetic control of DMA secretion. In our study, however, the treatment effect was stronger than the genotype effect. Zn deficiency stimulated DMA exudation in both genotypes, and although RIL46 showed higher absolute DMA release, IR74 displayed the larger relative fold-change (2.6 vs 1.9; Fig. 5).

This discrepancy between studies may reflect differences in environmental conditions (such as temperature and light intensity) and stress severity, which may have amplified treatment effects in our system.

#### 4.4.2 Comparative assessment of Zn accumulation and fold-change responses under Zn deficiency

The comparison of absolute Zn uptake under Zn-deficient conditions (Table 3) shows that RIL46 and IR74 perform similarly when Zn supply is limited. Across studies, the reported values are of comparable magnitude, with only modest differences between genotypes (e.g. 31 vs 18 µg plant^−1^ in Widodo et al.; 26 vs 18 µg g^−1^ in Arnold et al.; 14 vs 12 mg kg^−1^ in Rose et al.). Despite differences in units and experimental systems, the overall pattern is consistent. This suggests that the ability to uptake Zn under deficiency condition is not significantly different between the two genotypes.

The main difference between genotypes appears under Zn-sufficient conditions (+Zn), where IR74 consistently accumulates more Zn than RIL46 across studies (e.g., 212 vs 193 µg plant^−1^ in Widodo et al.; 189 vs 164 µg g^−1^ in Arnold et al.; 40 vs 28 mg kg^−1^ in Rose et al.). Because IR74 starts from a higher value under +Zn supply, its relative fold change and percentage reduction under Zn deficiency are correspondingly larger than those of RIL46 (Table 3; Fig. 6). Fig. 6 summarises genotype responses to Zn deficiency across studies. The heatmap (Fig. 6a) shows fold-change in Zn uptake (NoZn/+Zn with IR74 generally showing a larger reduction than RIL46, reflecting its higher Zn accumulation under Zn-sufficient conditions. The scatter plot (Fig. 6b) presents the percentage reduction in Zn uptake and shows that IR74 generally exhibits a greater relative decrease than RIL46 across studies. This does not necessarily imply that IR74 has lower capacity for Zn uptake under deficiency but rather reflects the contrast between its high uptake under Zn-sufficient conditions and its lower uptake under deficiency. Statistical comparisons between genotypes within the same treatment (Control RIL46 vs Control IR74; NoZn RIL46 vs NoZn IR74) showed no significant differences (P > 0.05), indicating that genotype alone does not strongly influence Zn accumulation under identical conditions of Zn supply.

Based on our data, differences in Zn uptake under deficiency conditions appeared to be driven primarily by Zn availability rather than genotype. The greater fold-change sensitivity observed in IR74 reflects its higher baseline uptake under Zn-sufficient conditions. For breeding and biofortification, tolerance traits likely need deeper physiological and molecular study beyond total Zn measurements. Fold-change trends provide useful indications, but further replicated research are needed to confirm these patterns.

Our results show that Zn deficiency strongly stimulates DMA exudation in all genotypes, including sensitive ones, confirming that DMA is part of the stress response. However, this increase does not translate into higher Zn uptake under deficiency conditions. These findings indicate that DMA exudation alone cannot fully explain tolerance to Zn deficiency.

## Acknowledgements

This project has received funding from the European Union’s Horizon 2020 research and innovation program under the Marie Sklodowska-Curie grant agreement No 101032337. Matthias Wissuwa has been partly funded by the Deutsche Forschungsgemeinschaft (DFG, German Research Foundation) under Germany’s Excellence Strategy – EXC 2070 – 390732324.

## Author contributions

C.R., G.L.M., C.T., M.W., R.V. and D.J.W. conceived the experiments, developed the methods and analysed the results. The experimental data were collected by C.R. assisted by G.L.M., C.T. and D.J.W. Data analysis and writing of paper by all authors.

## Data availability

The data sets generated and/or analysed during the current study are available on Zenodo, https://zenodo.org/uploads/18184803

