## Supplementary Information for "Significant increase in root exudation of 2′-deoxymugineic acid (DMA) as a response to zinc deficiency in rice"

| | | |
|---|---|---|
| 1 | A69-1 | **Tolerant** |
| 2 | RIL46 | **Tolerant** |
| 3 | IR55179 | **Tolerant** |
| 4 | IR64 | **Moderate** |
| 5 | IR74 | **Intolerant** |

**a)**

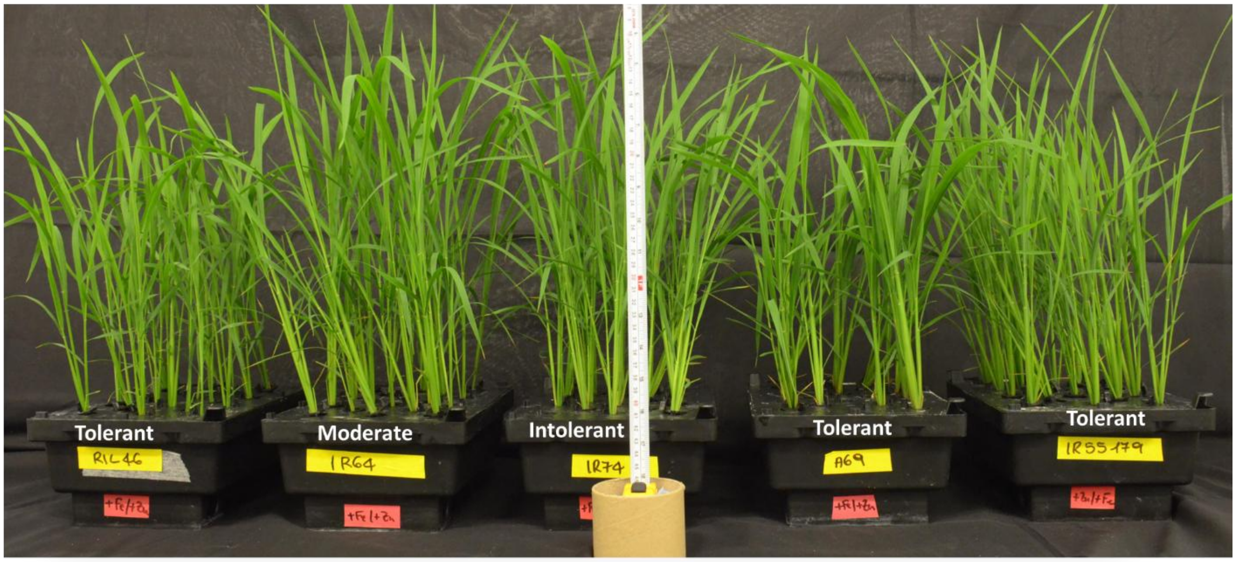


**b)**

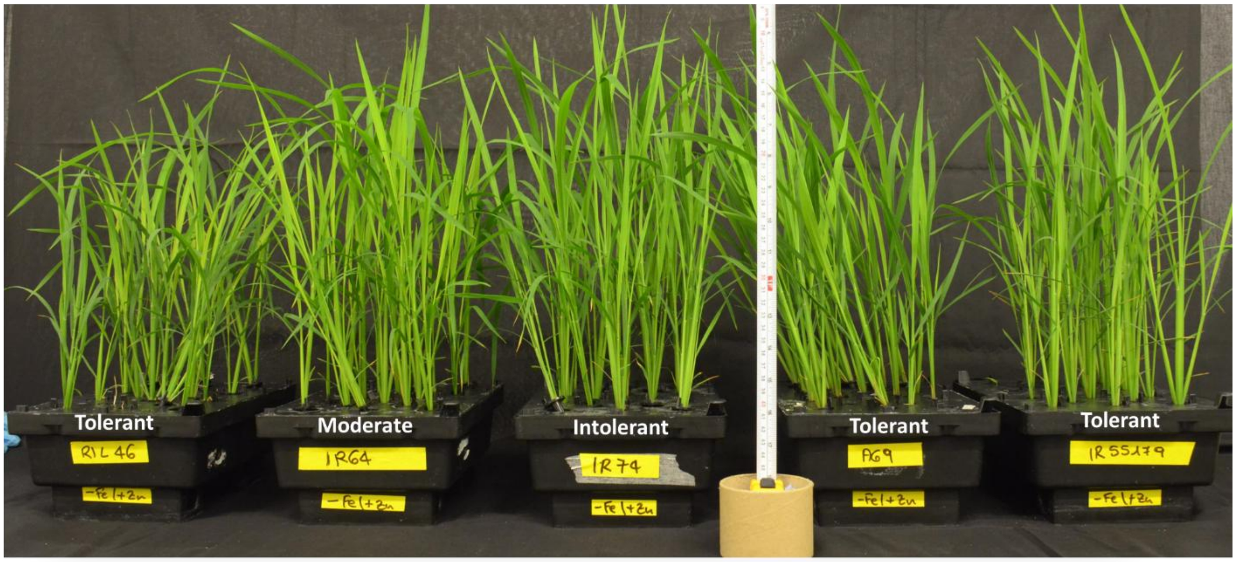

**c)**

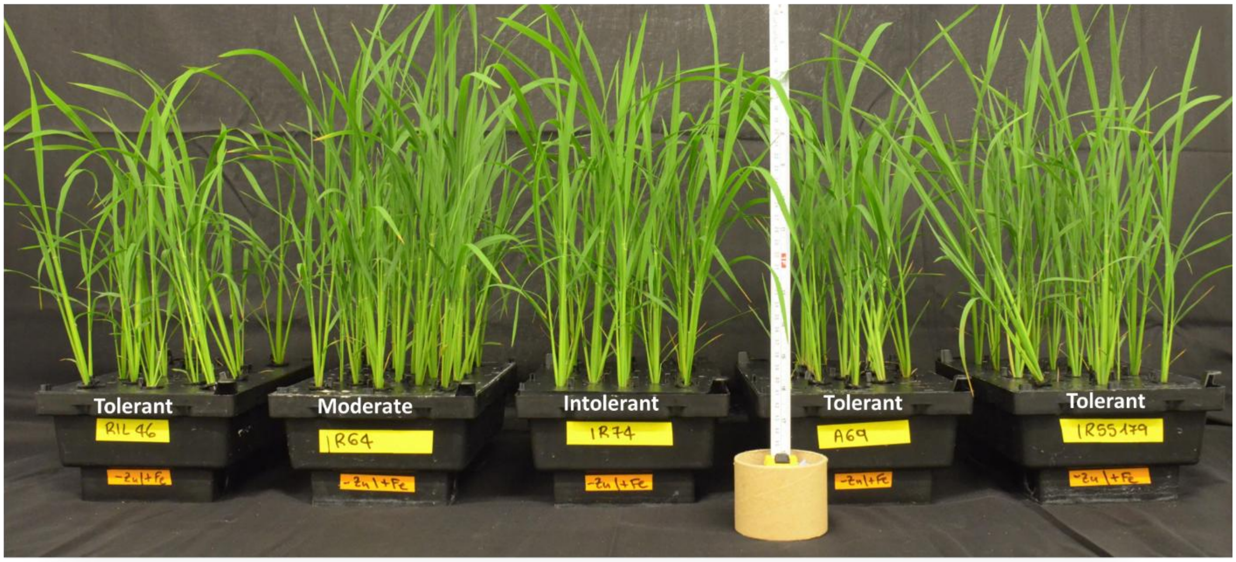


**Fig. S1** Photos of rice genotypes RIL46, IR64, IR74 A69-1, and IR55179 grown under (a) control conditions, (b) iron deficiency (-Fe), and (c) zinc deficiency (-Zn).

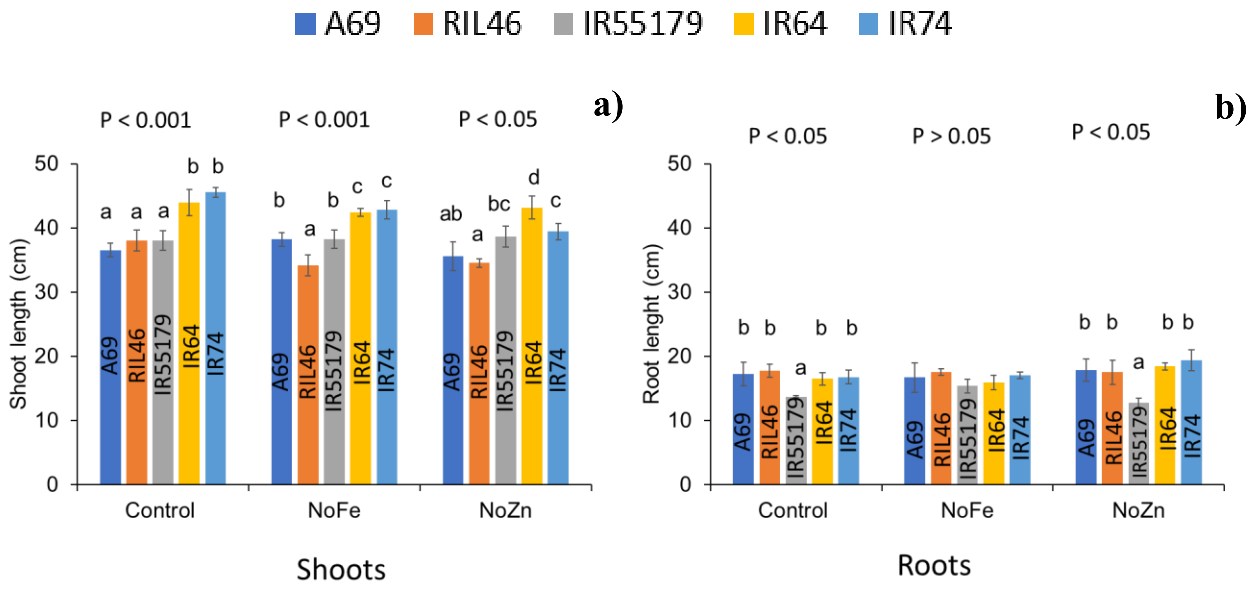


**Fig. S2** Shoot (a) and root (b) length, expressed in cm, of five different rice genotypes grown under iron (NoFe) and zinc (NoZn) deficiency treatments. Control treatment represents the complete nutrient solution. Values are presented as mean ± SD. Columns with different letters indicate statistically significant differences among genotype within the same treatment at $P < 0.05$.

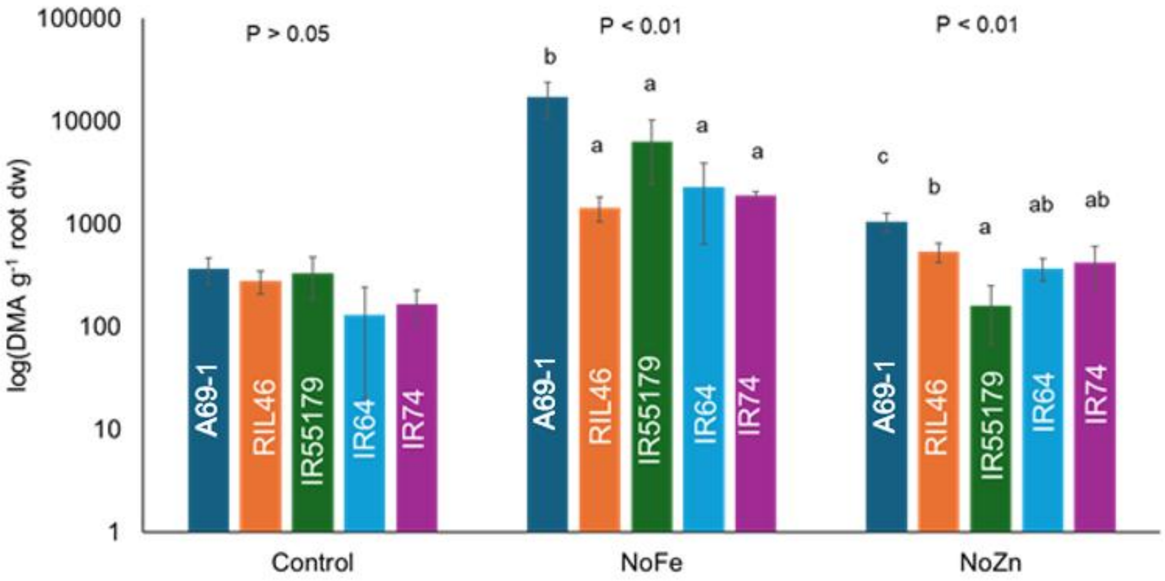


**Fig. S3** DMA exudation in different rice genotypes grown under iron and zinc deficiency conditions in hydroponic. Different letters indicate significant difference between means, based on ANOVA at $P < 0.01$

**Table S1** Iron (Fe) and zinc (Zn) concentrations (µM) in nutrient solutions under different treatments at week 1 and week 2.

Control represents the complete nutrient solution, while –Zn and –Fe indicates zinc- and iron-deficient treatments, respectively. Measurements were performed to ensure no contamination occurred and to confirm that –Zn and –Fe treatments effectively lacked Zn and Fe, respectively.

| | Sample Name | µM | |
|---|---|---|---|
| | | Fe | Zn |
| week 1 | control | 52.89 | 0.39 |
| | -Zn | 28.65 | 0.09 |
| | -Fe | 0.20 | 0.15 |
| Week 2 | control | 77.42 | 0.19 |
| | -Zn | 68.21 | 0.03 |
| | -Fe | 0.21 | 0.14 |

**Table S2** Observed m/z values of DMA in rice exudate processed in ESI- mode QTOF and relative abundance for the different nutrient deficiency treatments.

| Treatment | m/z | Abundance |
|---|---|---|
| Control | 303.1199 | 18563.96 |
| NoZn | 303.1198 | 15070.10 |
| NoFe | 303.1196 | 64768.50 |
